# Transient amplification enhances the persistence of tropicalising coral populations in marginal high latitude environments

**DOI:** 10.1101/2021.04.23.441133

**Authors:** James Cant, Katie Cook, James D. Reimer, Takuma Mezaki, Masako Nakamura, Cliodhna O’Flaherty, Roberto Salguero-Gómez, Maria Beger

## Abstract

Predicting the viability of species exposed to increasing climatic stress requires an appreciation for the mechanisms underpinning the success or failure of marginal populations. Rather than traditional metrics of long-term population performance, here we illustrate that short-term (*i*.*e*. transient) demographic characteristics, including measures of resistance, recovery, and compensation, are fundamental in the poleward range expansion of hard corals, facilitating the establishment of coral populations at higher-latitudes. Through the annual census of tropical and subtropical *Acropora* spp. colonies in Japan, between 2017-2019, we show how the transient amplification potential of a subtropical coral population supports its enhanced growth within unstable environmental conditions. The transient dynamics of both the tropical and subtropical populations were strongly influenced by their corresponding recruitment patterns. However, we demonstrate that variation in colony survival and fragmentation patterns between the two populations determines their relative capacities for transient amplification. This latitudinal variation in the transient dynamics of *Acropora* spp. populations emphasizes that coral populations can possess the demographic plasticity necessary for exploiting more variable, marginal conditions.

## Introduction

The latitudinal diversity gradient, or poleward decline in biodiversity (von Humboldt 1808), is a fundamental macroecological pattern evident across all major taxa (Hillebrand 2004; Fine 2015). This pattern emerges partly due to increased climatic variation at higher latitudes (Willig *et al*. 2003; Archibald *et al*. 2010; Mannion *et al*. 2014). Increased environmental variation exerts a strong filter on the assembly of biological communities, selecting for species with broader ecological niches (Janzen 1967). Yet, corresponding with the changing global climate, many ecosystems face imminent reassembly as species distributions shift to track favourable conditions (Pecl *et al*. 2017; Williams & Blois 2018). Along shifting distributional boundaries, the endurance of populations depends on their ability to withstand abiotic fluctuations (Valladares *et al*. 2014). Across a given species’ range, its populations are exposed to a series of environmental pressures giving rise to contrasting abilities between core and peripheral populations for tolerating abiotic variation (Angert 2009; Purves 2009). However, whilst the extent to which marginal populations can embrace environmental variation underpins the continued viability of numerous species, it is poorly understood how variation in the attributes that define the life cycles of species, such as longevity and age at reproduction, influences the persistence of populations along range boundaries (Valladares *et al*. 2014; Paniw *et al*. 2018; Healy *et al*. 2019).

Assessments of population viability typically explore long-term asymptotic dynamics, such as estimates of population growth rate (λ; Beissinger & Westphal 1998; Crone *et al*. 2011; Selwood *et al*. 2015). However, evaluating the transient, or short-term, dynamics of natural populations is as important, if not more so, for anticipating the persistence of various species (Hastings 2001, 2004; McDonald *et al*. 2016). The transient dynamics of populations reflect their dynamics within unstable environments, describing how a population’s trajectory can change in the short-term relative to its asymptotic growth rate (Table 1; Stott *et al*. 2011).

**Table 1.**
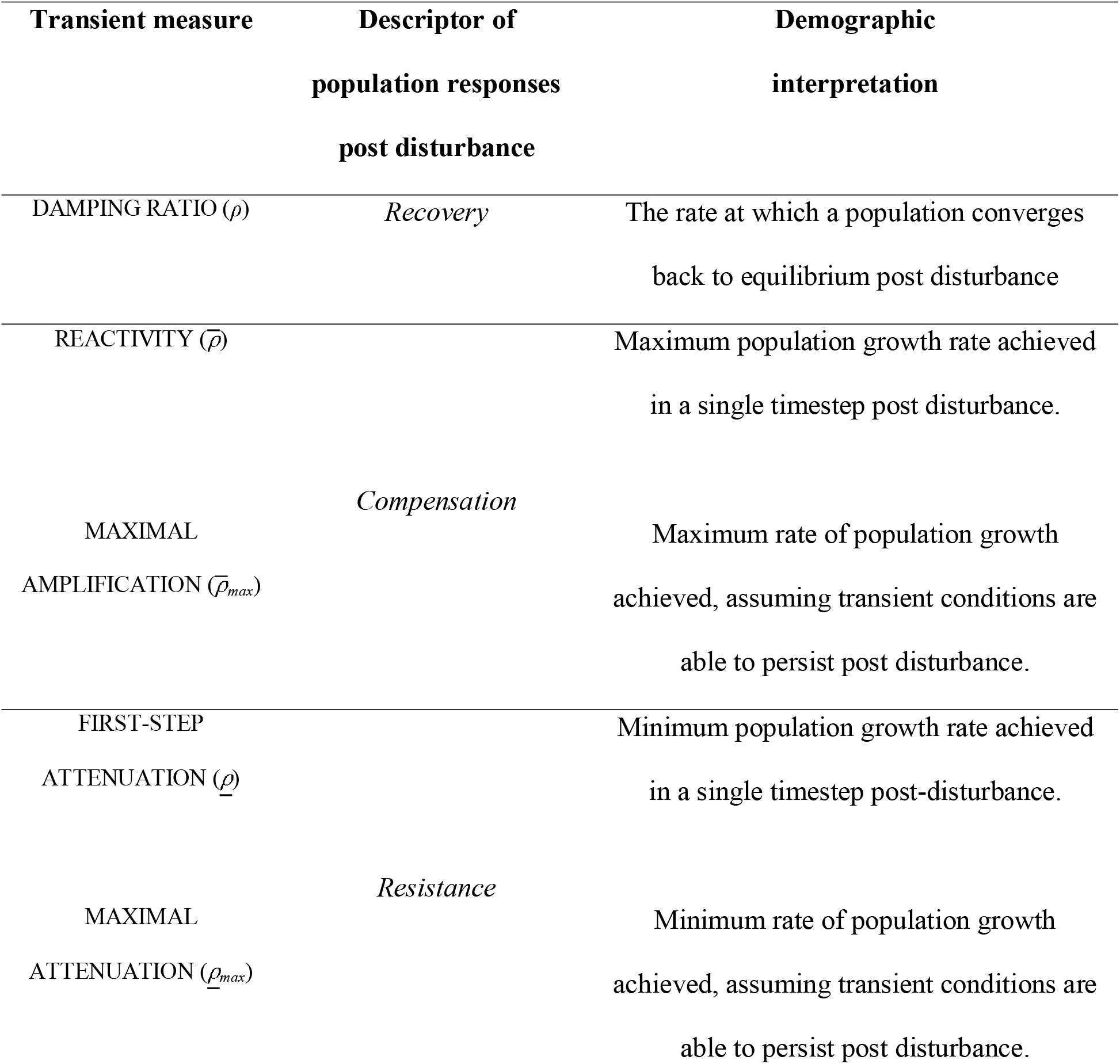
Summary of the metrics used here to describe the transient (short-term) dynamics of tropical and subtropical *Acropora* spp. populations, alongside their corresponding disturbance response descriptor (resistance, compensation, and recovery) and demographic interpretation.

Transient dynamics therefore provide a convenient means for quantifying population resilience, specifically, the ability of populations to resist and recover after disturbances (Capdevila *et al*. 2020). Driven by rapidly changing climate regimes and intense anthropogenic pressure, many ecosystems are at risk of bifurcation, *i*.*e*. the loss of an equilibrium state (*sensu* Poincaré 1885). Following a bifurcation, transient dynamics can provoke the increase (*amplification*) or decline (*attenuation*) of a population. Thus, understanding and predicting the transient dynamics of populations has become a priority for pest management and conservation (Ezard *et al*. 2010; Hodgson *et al*. 2015; Capdevila *et al*. 2020).

Global warming, together with strengthening poleward boundary currents, are driving the rapid tropicalisation of marine communities along tropical to temperate transition zones (Vergés *et al*. 2014; Kumagai *et al*. 2018). Consequently, tropical taxa, including many zooxanthellate hard coral species, are becoming increasingly prevalent in high-latitude subtropical environments (Denis *et al*. 2013; Vergés *et al*. 2019). This establishment of coral populations along subtropical coastlines draws many similarities with the dynamics of biological invasions, which, economic and ecological costs aside, represent the growth of small populations within novel environments (Iles *et al*. 2016). Particularly relevant in this context is evidence that the transient dynamics of plant populations are effective predictors of invasive potential (Iles *et al*. 2016). Indeed, populations possessing the capacity for rapid amplification following a perturbation (reflected here by the introduction of a novel environment) are more capable at exploiting new habitats (Jelbert *et al*. 2019). It can be expected, therefore, that the capacity of coral populations for establishing at higher latitudes may be dictated by their transient dynamics, rather than asymptotic population trajectories. Nevertheless, the transient dynamics of coral populations remain unexplored (Cant *et al*. 2021a).

Here, we explore if and how variation in the transient dynamics of coral populations is consistent with their exposure to abiotic variability. Specifically, we compare the relative stability (*attenuation* and *amplification;* see Table 1) and recovery attributes of tropical and subtropical *Acropora* spp. populations in southern Japan; a region considered an epicentre of tropicalisation (Vergés *et al*. 2014; Kumagai *et al*. 2018). Transient dynamics are thought to buffer the effects of environmental variability and are therefore accentuated in populations exposed to more frequent disturbances (Ellis & Crone 2013). Accordingly, we investigate whether coral populations at higher latitudes exhibit more pronounced transient dynamics than their tropical counterparts. Equally, the reproductive isolation associated with high-latitude coral populations ensures that they are typically supported by sporadic recruitment from up-current tropical reefs, with their endurance instead reliant on the dynamics of existing colonies (Cant *et al*. 2021b). Subsequently, we also conduct a transient Life Table Response Experiment (Koons *et al*. 2016) decomposing variation in the transient dynamics of tropical *vs*. subtropical *Acropora* spp. populations to test whether the transient dynamics of subtropical coral populations are indeed sustained by the dynamics of existing colonies.

## Materials and Methods

### Model parameterisation

To explore the influence of environmental variability on the transient dynamics of coral populations, we utilised an Integral Projection Model (IPM) framework (Easterling *et al*. 2000) to quantify the respective dynamics of *Acropora* spp. populations from a tropical and subtropical environment. An IPM describes how size-specific vital rates (e.g., survival, recruitment) observed at the individual-level translate into population-level characteristics:

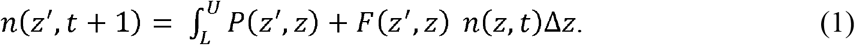

Here, the size (*z*; in this case, colony surface area, cm^2^) structure, *n*(*z*’, *t* +*1*), of a population at time *t*+*1* is a function of its structure at time *t, n*(*z, t*), and the demographic patterns outlined by the sub-kernels *P* and *F. P* describes size-specific patterns relating to colony survival probability (σ), transitions in size (*γ*; growth, stasis, and shrinkage), the probability of fragmentation (*κ*), and the number and size of fragments produced (*κ*_*n*_ and *κ*_*0*_, respectively):

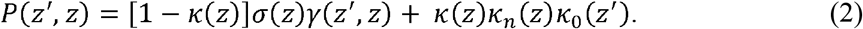

*F* describes the recruitment of new, sexually produced individuals (*C*_*0*_), which are the outcome of colony fecundity (larval production per colony, *f*_*n*_), constrained by both the probabilities of larval settlement (*ψ*), and post-settlement survival (⍰):

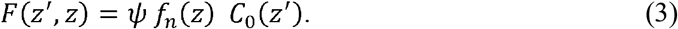

All size-specific vital rates reflect patterns estimated across a size range (Δ*z*) equal to 10% above and below the maximum (*U*) and minimum (*L*) observed sizes for the studied populations to avoid accidental eviction (Williams *et al*. 2012).

We empirically parameterised our IPMs through the annual census of tropical and subtropical *Acropora* spp. populations in southern Japan (Fig. 1). The *in situ* identification of *Acropora* colonies to species level is complicated by the widespread occurrence of morphologically cryptic subspecies and species hybridisation (Richards & Hobbs 2015; Richards *et al*. 2016). Thus, working at the genus-level we pooled data from across repeated surveys of tagged individuals in September 2017, August 2018, and August 2019, to calculate region- and size-specific patterns in colony survival (σ), transitions in size (*γ*), and fragmentation (*κ*; Supplementary S1). Colony survival represented the continued presence of tagged colonies over time and was modelled as a function of colony size at time *t*. Alternatively, transitions in colony size reflected the difference between colony surface areas recorded during successive annual surveys. In this context, transitions in colony size reflected both growth due to colony extension, and shrinkage following partial mortality (Madin *et al*. 2020), and was calculated using the relationship between colony size at time *t* and at time *t*+*1*. Next, using data pooled from both the tropical and subtropical populations, we modelled the probability of colony fragmentation as a function of colony size at time *t*. This approach was necessary due to the low frequency of annual fragmentation events (number of events reported, 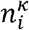) observed within our tropical population, although we weighted fragmentation probabilities according to the relative proportion of annual events recorded across the tropical and subtropical populations (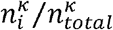; see Supplementary S1 for further details). Finally, we estimated patterns in fragment production (*κ*_*n*_) and fragment size (*κ*_*0*_) as a function of initial colony size, using the number and recorded size of all observed colony fragments.

**Figure 1.**
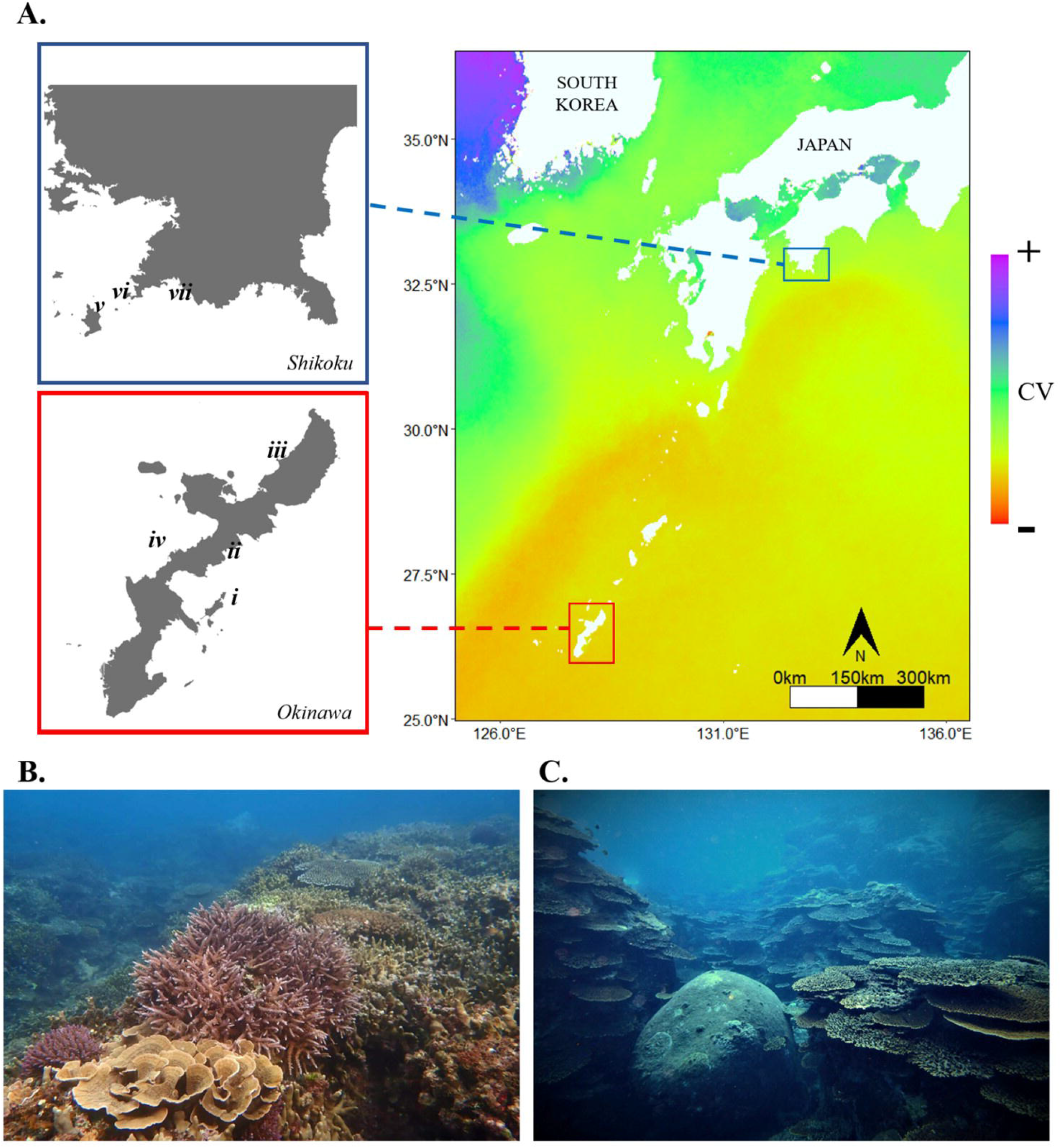
The locations of the surveyed tropical and subtropical *Acropora* spp. populations in Japan, separated by a distance of 990km. **(A)** Mediated by the Kuroshio Current, the coastline of southern Japan aligns with a distinct gradient in environmental variability (coefficient of variation, CV) in monthly sea surface temperatures (SSTs) recorded during our sampling years between 2017 and 2019. We tagged individual *Acropora* spp. colonies at four locations within the tropical reef communities of Okinawa (Red): (i) Miyagi Channel, (ii) Oura Bay, (iii) Hentona, and (iv) Onna (only visited for deploying settlement tiles used to quantify recruitment patterns), and at three locations within the subtropical coral communities of Kochi (Blue): (v) Okinoshima, (vi) Kashiwajima, and (vii) Nishidomari. Representative photographs of surveyed tropical and subtropical coral assemblages at **(B)** Hentona, Okinawa, and **(C)** Kashiwajima, Kochi. Photo credits: K. Cook.

In our IPMs, recruitment encompassed patterns in colony fecundity (*f*_*n*_), and the probabilities of larval settlement (*ψ*) and post-settlement survival (henceforth recruit survival probability [*⍰*]). Although we did not directly measure colony fecundity due to the logistical challenges involved (Gilmour *et al*. 2016), we estimated annual larval output (volume of larvae produced, cm^3^) as a function of colony size using a relationship reported in *Acropora* spp. on the Great Barrier Reef (Supplementary S2; Hall & Hughes 1996). Additionally, we determined the probabilities of larval settlement and recruit survival, using larval counts made during prior tropical (2011–13; Nakamura *et al*. 2015) and subtropical (2016–18; Nakamura, *unpublished data*) settlement tile surveys in southern Japan (see Supplementary S2 for further details). Combining the larval counts per unit area from these earlier surveys with our regional estimates of larval output and observed recruit densities, enabled us to estimate ratios translating colony larval output from a measure of larval volume into expected counts of settling larvae (*ψ*; *sensu* Bramanti *et al*. 2015), and define a series of post-settlement survival probabilities reflecting temporal trends in recruitment success within both a tropical and subtropical setting (*⍰*; Supplementary S2). Finally, consistent with evidence that the dynamics of larval settlement and survival are strictly coordinated by synergistic abiotic constraints (Vermeij *et al*. 2009; Doropoulos *et al*. 2016), we modelled the size distribution (*C*_*0*_) of tropical and subtropical recruits independently of parent colony size.

### Quantifying transient dynamics

We used our IPMs to test our hypothesis of variation in the transient dynamics of tropical *vs*. subtropical coral populations. We focused on transient measures depicting the demographic resilience attributes described in Table 1: recovery (*damping ratio* [*ρ*]), resistance (*first-step attenuation* [*<u>ρ</u>*] & *maximal attenuation* [*<u>ρ</u> max*]), and compensation (*reactivity* 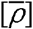 & *maximal amplification* 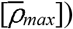). To obtain estimates of variance in these transient metrics, we generated 1,000 variants of our tropical and subtropical IPMs using Jack-knife resampling; each time omitting 5% of our data without replacement whilst allowing the modelled probabilities of larval settlement (*ψ*) and recruit survival (*⍰*) to vary within observed limits. Next, we integrated the kernel of each model variant into a high-dimension matrix (200×200 cells) using the ‘midpoint rule’ (Ellner & Rees 2006; Zuidema *et al*. 2010), with the probability of individuals transitioning from one cell to the next estimated at the cell midpoint and multiplied by the cell width. In our case cell width corresponded with colony size increments of 0.716 cm^2^ on the log-scale. Following this discretisation, we calculated the distribution (mean and variance) of each transient metric for the tropical and subtropical populations using the R package *popdemo* (Stott *et al*. 2012).

We calculated the amplification and attenuation characteristics of the tropical and subtropical populations as population structure-specific measures. Population structure-specific transient measures provide the predicted transient dynamics of a population given its current state distribution; as opposed to transient bounds which reflect the potential dynamics of a population irrespective of its state distribution (Stott *et al*. 2011). For these calculations, we derived the state distributions of both the tropical and subtropical *Acropora* spp. populations using the size distributions of tagged colonies recorded during our 2019 census. Across our Jack-knife model variants, some combinations of resampled vital rate schedules lacked the capacity for eliciting either amplification or attenuation in their corresponding population relative to asymptotic growth rates. We therefore present the percentage of model variants from which predictions of amplification and attenuation could be obtained as an additional indication of the transient potential of the tropical and subtropical *Acropora* spp. populations. Finally, to contextualise our estimates of transient dynamics against the long-term trends of each population, we calculated mean and variance estimates of their asymptotic growth rates (λ), with λ < 1 or > 1 reflecting negative or positive population growth (Caswell 2001).

### Model decomposition

We tested our hypothesis that the transient dynamics of subtropical coral populations are sustained by the survival, transitions in size, and fragmentation patterns of existing colonies rather than patterns in recruitment using a transient Life Table Response Experiment (transient LTRE; Koons *et al*. 2016). The amplification characteristics of populations define their capacity to exploit and thrive within novel, variable environments (McDonald *et al*. 2016; Jelbert *et al*. 2019). Thus, we decomposed the vital rate influences of the relative maximal amplification characteristics 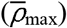 of tropical *vs*. subtropical *Acropora* populations. The transient dynamics of our focal coral populations (*ξ*) are contingent on three components: the size-specific vital rate patterns of established colonies (*Θ*), and the probabilities of larval settlement (*ψ*) and recruit survival (*⍰*). Variation in these components between the tropical and subtropical populations consequently drives any variation between the characteristics of the two populations:

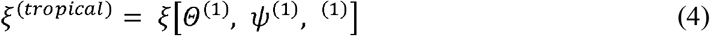

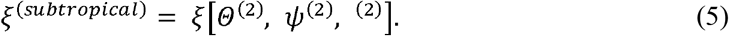

Within coral populations, rates of larval settlement and survival oscillate considerably over time (Davidson *et al*. 2019). Thus, we incorporated this variability into our IPMs by allowing the probabilities of larval settlement and recruit survival to fluctuate within observed boundaries, therefore introducing an element of within–population variability to our models. Using the transient LTRE approach detailed below, we combined a traditional Life Table Response Experiment with a Kitagawa & Keyfitz decomposition (Kitagawa 1955; Keyfitz 1968; Caswell 2019). Briefly, this decomposition approach allowed us to account for within-population variability when evaluating the vital rate mechanisms underlying the differences between the transient dynamics of two populations (Maldonado-Chaparro *et al*. 2018; Layton-Matthews *et al*. 2021).

We first paired up tropical and subtropical model variants to evaluate the overall contributions (*C*) of the vital rate patterns of established colonies (*Θ*), larval settlement (*ψ*), and recruit survival (*⍰*), towards variation in 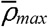 using a Kitagawa & Keyfitz decomposition. The overall contribution of each component was obtained by averaging the effect on 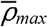 of substituting the tropical and subtropical form of the selected component against a fixed background of the other components (Caswell 2019):

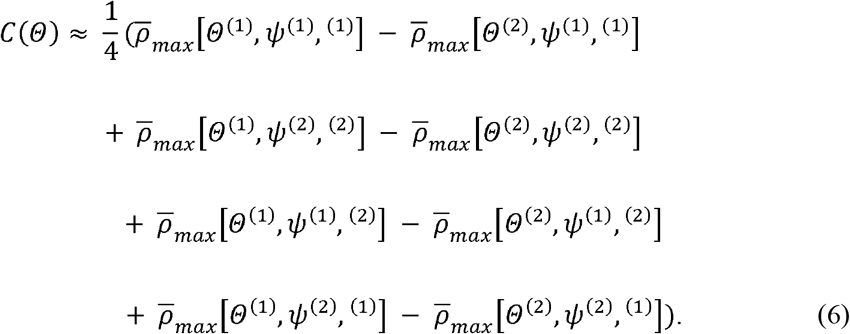

Across all decomposition analyses, we defined the tropical population as our control model. Subsequently, positive contributions reflect greater influence towards the dynamics of the tropical population, whereas negative contributions imply a greater importance towards the subtropical population.

Next, we decomposed the separate contributions of the vital rates of survival, changes in size, and fragmentation, observed in established colonies, towards variation in 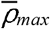. The contribution of each vital rate (C[*θ*_*i*_]) corresponds with the change in that vital rate between paired tropical and subtropical models combined with the environmental-specific elasticity matrices of 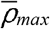 (Caswell 2019):

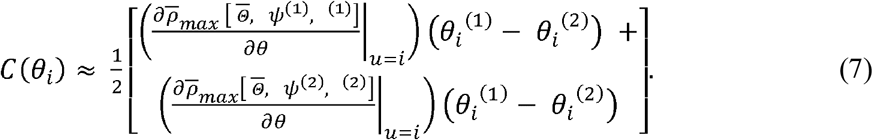

Here, the tropical- and subtropical-specific elasticity matrices of 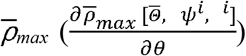 were comprised of the proportional sensitivities (*e*_*ij*_) of 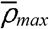 towards the matrix elements (*a*_*ij*_) of a discretised IPM kernel parameterised using the mean vital rates across our tropical and subtropical populations 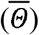:

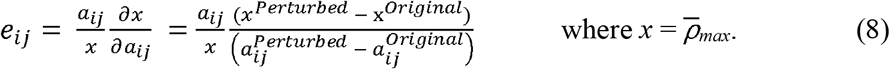

## Results

### Trends in transient dynamics: Tropical vs. subtropical

The transient characteristics of the subtropical *Acropora* spp. population were more pronounced than in its tropical counterpart (Fig. 2). Of the two populations, the tropical *Acropora* spp. population displayed the highest asymptotic growth rate (λ: Tropical = 0.916 [95% CI: 0.914, 0.918]; Subtropical = 0.655 [0.654, 0.655]). However, the subtropical population is expected to exhibit a reactive 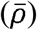 transient response to perturbation, experiencing an increase in its growth rate relative to its asymptotic trajectory 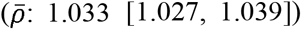. Although, across all Jack-knifed model variants, the subtropical variants presented more heterogenous responses to perturbations, than the tropical variants (Fig. 2A). Alternatively, post disturbance, the tropical population is predisposed to experience attenuation (*<u>p</u>*), resulting in a decline in its population growth rate (*<u>p</u>*: 0.985 [0.983, 0.986]).

**Figure 2.**
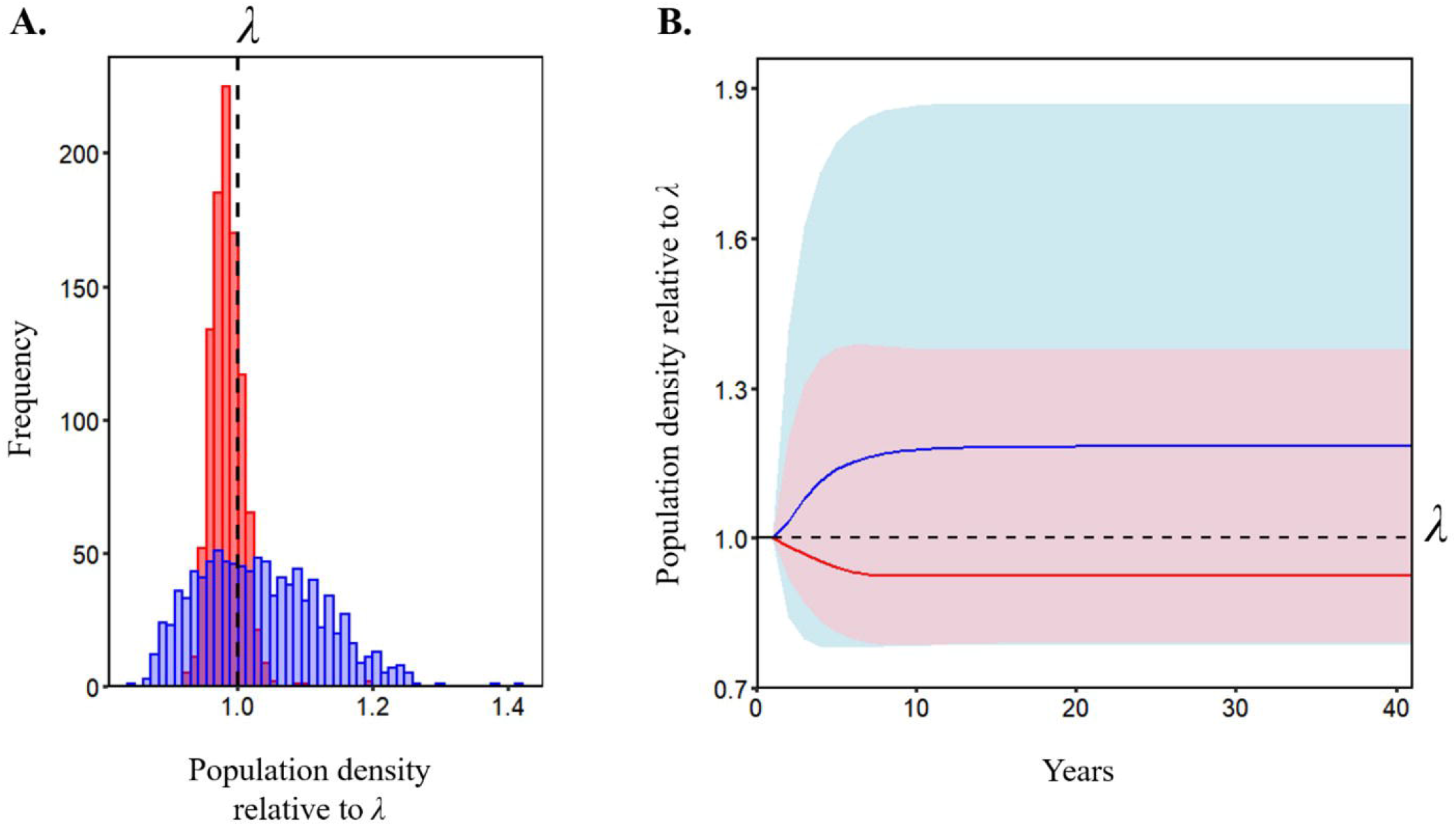
The subtropical *Acropora* spp. population has an enhanced capacity for demographic amplification compared to its tropical counterpart. Using Jack-knife resampling we generated 1,000 Integral Projection Model variants for the subtropical and tropical *Acropora* spp. populations, each time omitting 5% of surveyed colonies whilst allowing larval settlement and recruit survival probabilities to vary randomly within observed limits. From each model variant we estimated measures of transient (short-term) dynamics describing the dynamics of the tropical (Red) and subtropical (Blue) *Acropora* spp. populations following disturbance. **(A)** The distribution of transient responses within one time–step of a perturbation, observed across the model variants. **(B)** Illustrates how the transient dynamics of the model variants manifest over 40 years post-disturbance modifying population trajectories relative to original asymptotic expectations. Solid lines represent the mean population trends with shaded areas reflecting the range of observed transient patterns for each population. Across both panels all transient responses in population size are displayed relative to each population’s corresponding asymptotic population growth rate (λ, dashed line).

Notably, in comparison with the tropical population, the transient dynamics of the subtropical population demonstrate a superior capacity for maintaining elevated population growth within unstable environments (Fig. 2B). Amplification was observed in 84.5% of subtropical model variants as opposed to in just 23.1% of tropical variants. Indeed, expected maximal amplification 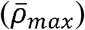 was also highest within the subtropical population, and reflected a potential ∼22% increase in population growth rate following a disturbance relative to asymptotic expectations (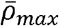. Tropical = 1.019 [1.012, 1.026]; Subtropical = 1.228 [1.215, 1.241]). The tropical population did display a higher damping ratio (*ρ*) than the subtropical population (*ρ*. Tropical = 1.638 [1.634, 1.641]; Subtropical = 1.429 [1.424, 1.433]), indicating a faster convergence rate to an equilibrium state. However, in this context, this disparity in convergence rate corresponds with the more prominent transient displacement observed in the subtropical model variants relative to their asymptotic characteristics (Fig. 2B). Conversely, maximal attenuation(*<u>ρ</u>*_*max*_) estimates for the tropical and subtropical populations suggest that, whilst attenuation was more readily observed within tropical model variants (observed in 96% and 40.3% of tropical and subtropical variants respectively), both populations are only expected to experience a <10% reduction in their growth rates relative to asymptotic expectations should attenuation occur (*<u>ρ</u>*_*max*_. Tropical = 0.919 [0.916, 0.923]; Subtropical = 0.940 [0.935, 0.946]).

### Transient LTRE decomposition

Despite clear evidence that recruitment patterns shape the transient dynamics of the tropical and subtropical *Acropora* spp. populations, the differential vital rate schedules of existing colonies are responsible for the variation observed between the amplification capacities of the two populations (Fig 3). Patterns in larval settlement (*ψ*), recruit survival (*⍰*), and the vital rates of existing colonies (*Θ*) varied significantly in their contributions towards variation in the maximal amplification 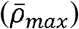 estimates obtained for the tropical and subtropical populations (ANOVA: F_2, 2997_ = 29557, *p* < *0*.*001*; Tukey: *ψ* > *Θ* > *⍰*). Overall, larval settlement (*ψ*) and recruit survival (*⍰*) exerted the greatest influence on estimates of 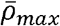, although the influence of these two properties was not consistent across the tropical and subtropical populations (Fig. 3A). The amplification characteristics of the subtropical population were underpinned by patterns in larval settlement, whereas the corresponding characteristics in the tropical population were guided by patterns in recruit survival (see Supplementary S2). Ultimately, these contrasting trends served to nullify the proportional contribution of recruitment dynamics towards variation in 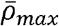 between the tropical and subtropical populations.

**Figure 3.**
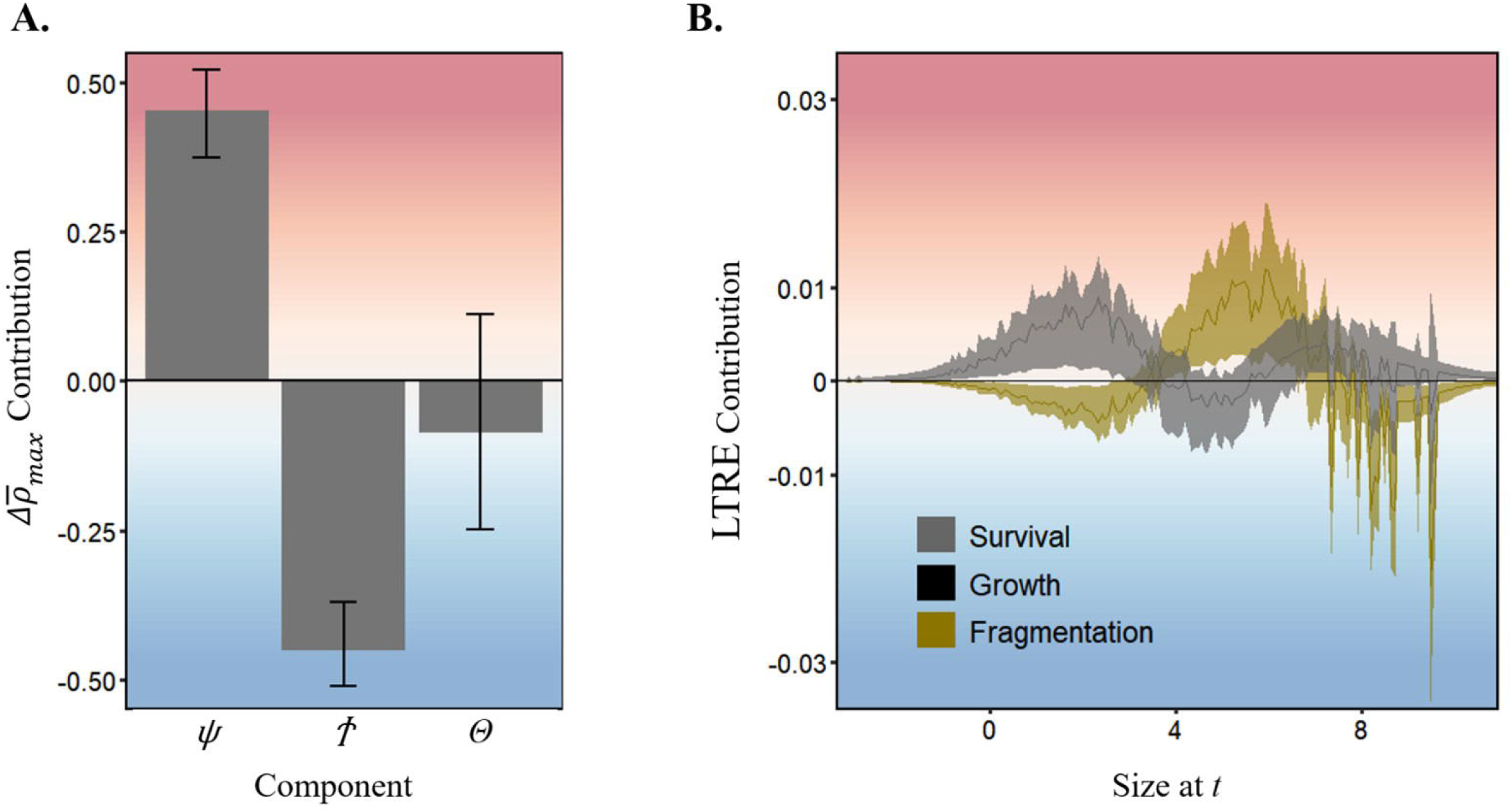
Size-specific patterns in colony survival and fragmentation underpin the varying amplification characteristics of the tropical and subtropical *Acropora* spp. populations. **(A)** The proportional contribution of patterns in larval settlement (*ψ*), recruit survival (*⍰*), and the vital rate schedules of existing colonies (*Θ*) towards differences in the maximal amplification characteristics of tropical and subtropical *Acropora* spp. populations 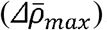. **(B)** Relative size-specific contributions of the vital rates of survival, growth, and fragmentation towards differences between the maximal amplification characteristics of the subtropical *Acropora* spp. population compared with its tropical counterpart as a baseline. Solid lines represent mean contribution patterns. Across both panels, positive contributions reflect greater influence of a given vital rate towards the transient characteristic reported for the tropical population (Red), whereas negative values reflect greater influence towards the subtropical population (Blue). All error displayed represents the full range of observations observed across tropical and subtropical model variants.

Consequently, disparity in the dynamics of existing tropical and subtropical colonies, specifically their survival and fragmentation characteristics, underpinned the contrasting amplification capacities of their corresponding populations (Fig. 3B). Although variable, the relative contribution of colony survival towards variation in 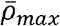 decreased with colony size, such that regional variation in the survival patterns of small colonies (0.37 to 55 cm^2^; -1 to 4 cm^2^ on the log scale) strongly influenced estimates of 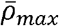 (Fig 3B). Alternatively, the contribution of colony fragmentation towards variation in 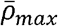 increased with colony size, with the fragmentation patterns of large (>1097 cm^2^, >7 cm^2^ on the log scale) subtropical colonies serving to enhance the amplification capacity of their population (Fig 3B). By contrast, colony growth characteristics did not influence the transient amplification potential of either population (Fig. 3B). Evidently, the enhanced amplification capacity of the subtropical coral population is associated with the fragmentation characteristics of larger colonies. Meanwhile, the elevated survival of smaller tropical colonies, relative to subtropical colonies, serves to diminish the amplification potential of the tropical population.

## Discussion

Global climatic change is reassembling coral reef communities worldwide (Hughes *et al*. 2017, 2018). Accordingly, understanding the mechanisms underpinning the establishment and persistence of range-shifting coral species in subtropical and temperate locations is imperative for anticipating the future success or failure of global coral assemblages, and their continued provision of essential ecosystem services (Hoegh-Guldberg *et al*. 2017; Camp *et al*. 2018; Sommer *et al*. 2018). Comparing between the dynamics of a tropical and subtropical *Acropora* spp. population we documented higher asymptotic population growth in the tropics. However, we have illustrated that the expansion and endurance of a coral population in a highly variable and, comparatively stressful, subtropical environment corresponds with its superior capacity for amplified population growth following disturbance compared to a down current tropical population (Fig. 2). We also discovered how the transient dynamics of a subtropical *Acropora* spp. population are contingent on the survival and fragmentation dynamics of existing colonies, highlighting key drivers underpinning the fitness of coral populations at higher latitudes. Recruitment had the largest overall effect on the dynamics of both the tropical and subtropical *Acropora* spp. populations. Yet, divergent larval settlement and recruit survival probabilities between the two populations ensures that the dynamics of existing colonies underpin the relative differences between their transient dynamics (Fig. 3). Overall, our findings here, are consistent with insights from invasive populations whose transient demographic characteristics facilitate the colonisation of non-native environments (Iles *et al*. 2016; Jelbert *et al*. 2019), evidencing mechanisms that shape the ability for coral species to shift their distributions into subtropical and temperate environments.

### Transient versus asymptotic dynamics

Understanding within-species demographic variation across climatic gradients is essential for forecasting the success of populations at tracking favourable conditions and establishing themselves within novel environments (Merow *et al*. 2017). Our findings display an emergent latitudinal trade-off between the long-term viability and short-term exploitation potential of *Acropora* spp. populations in southern Japan. Similar divergent latitudinal patterns in stability and variability have been observed across various biological scales (Hillebrand *et al*. 2018; Antão *et al*. 2020), and are thought to underpin the vulnerability of lower-latitude populations to future climatic change (Barlow *et al*. 2018). Across tropical and subtropical *Acropora* spp. populations, asymptotic population growth was highest in the tropics, aligning with traditional expectations that population growth rates will decline towards species range boundaries as populations encounter increasingly demanding environments (Vucetich & Waite 2003). However, the strength and universal nature of this expectation is widely refuted (Sagarin & Gaines 2002; Sexton *et al*. 2009; Villellas *et al*. 2013). Instead, peripheral populations have been demonstrated to exhibit greater temporal variability in population growth rates (Villellas *et al*. 2013). Indeed, maximising transient amplification potential is considered a more beneficial strategy than prioritising long-term population growth, for enhancing population persistence within unstable, marginal, environments (McDonald *et al*. 2016). Thus, whilst the tropical *Acropora* spp. population appears more viable under stable conditions, the subtropical population displays demographic strategies associated with the enhanced exploitation of more variable environments.

Peripheral populations inhabiting sub-optimal or more varied environments compared to core populations, are becoming increasingly crucial for species persistence under climate change (Valladares *et al*. 2014). The mechanisms behind the long-term viability of coral populations at higher latitudes have long been disputed (Beger *et al*. 2014). At higher-latitudes coral populations are susceptible to erosion (Nozawa *et al*. 2008), thermal stress (Kim *et al*. 2019; Cant *et al*. 2021b), reproductive and genetic isolation (Thomas *et al*. 2017; Precoda *et al*. 2018; Nakabayashi *et al*. 2019), and are exposed to cooler, highly seasonal abiotic regimes, and reduced irradiance (Yamano *et al*. 2012; Muir *et al*. 2015; Sommer *et al*. 2017). However, legacies of exposure to variable environments affords populations with greater adaptive capacity, as abiotic variability cultivates, and filters, the traits necessary for the tolerance of further disturbances (Kroeker *et al*. 2020). In subtropical coral communities, the maintenance of diverse gene pools largely relies on their connectivity with upcurrent tropical reefs, a characteristic that is restricted in many of these systems (Noreen *et al*. 2009; Beger *et al*. 2014). However, sporadic larval supply into subtropical coral communities may benefit their adaptation to abiotic variability, preventing genetic swamping from tropical ecosystems that experience radically different selection pressures (Galipaud & Kokko 2020). However, as marginal populations become increasingly fragmented or isolated, their diminished genetic diversity inhibits their durability within variable environments (Pearson *et al*. 2009). Thus, although we have demonstrated that coral populations display the demographic plasticity necessary for exploiting more variable regimes, the continued success of high-latitude coral populations is likely contingent on continued support from core populations (Cant *et al*. 2021b).

We note here that we conducted our demographic assessment of *Acropora* spp. populations at the genus level. Considering the prevalence of uncertainties within coral taxonomy (Fukami *et al*. 2004), there is a precedence for assessments into the characteristics of coral populations to operate at higher taxonomic levels (Darling *et al*. 2019; Edmunds 2020). Our interpretation of demographic plasticity therefore assumes a consistency in species configurations across our tropical and subtropical *Acropora* spp. populations. However, species records from both Okinawa and Kochi (see Nishihira & Veron [1995] and Veron *et al*. [2016]) indicate there is considerable overlap in the composition of these tropical and subtropical *Acropora* spp. populations (Supplementary S3). Equally, there is minimal variation in the morphological and functional traits of acroporid species associated with the coastal communities of Okinawa and Kochi (Supplementary S3), reinforcing our interpretations of the observed demographic variation between the two populations.

### Decomposing latitudinal contrasts within vital rate patterns

The size structure of coral populations has considerable repercussions on their dynamics and interactions within their wider reef communities (Dietzel *et al*. 2020; Pisapia *et al*. 2020). The heightened amplification characteristics we observed in the subtropical *Acropora* spp. population were primarily supported by the survival, and fragmentation patterns of larger individuals (Fig. 3). This pattern reflects our expectation that, with subtropical coral populations reliant on sporadic recruitment events, their endurance is conditional on the vital rates of existing colonies. Colony fragmentation is commonly observed within disturbed environments (Pisapia *et al*. 2019), and is a common trait amongst *Acroporid* species enabling the rapid colonisation of available substrate (Roth *et al*. 2013). Indeed, the growth of colony remnants following fragmentation has been shown to support faster rates of recovery in coral cover than the growth of recruits and younger colonies of equal size (Connell 1997). Along tropicalising coastlines, colonisation through individual fragmentation could prove particularly effective, with rising temperatures and grazing tropical migrants reducing macroalgal competition (Vergés *et al*. 2016; Kumagai *et al*. 2018), and limited accretion reducing the density of existing coral communities (Kleypas *et al*. 1999). However, increased fragmentation also implies an accumulation of smaller sized colonies, and is attributed with the diminishing capacity for coral populations to persist during recurrent climatic disturbances (Riegl *et al*. 2012; Riegl & Purkis 2015; Pisapia *et al*. 2019).

It is not unusual for the dynamics of coral communities to revolve around the vital rates of the largest colonies (Dietzel *et al*. 2020), yet the reliance of the subtropical *Acropora* spp. population on the dynamics of larger individuals could render it sensitive to future climate shifts. In Japan, the frequency of severe typhoon storms is increasing (Hoshino *et al*. 2016). These storms are known to disproportionally impact upon the largest individuals within coral communities, particularly those with delicate tabular and branching structures such as *Acropora* spp. (Bries *et al*. 2004; Madin & Connolly 2006). During September 2018, Typhoon Jebi, possessing wind speeds upwards of 158km/h, made landfall along the southern coastline of Shikoku Island (Mori *et al*. 2019). This storm exerted considerable structural damage within the subtropical coral communities of Kochi (Cant, Cook & Reimer, 2019, *pers. obs*.), and is deemed responsible for a decline in mean colony size we observed within the subtropical *Acropora* spp. population during 2019 (Supplementary S1). The dominance of larger sized colonies in this subtropical population (Supplementary S1) suggests that this population has successfully navigated past typhoon storms. However, with the intensity of future storms increasing (Hoshino *et al*. 2016), destructive events on the scale of Typhoon Jebi will become more frequent, possibly undermining the success of coral populations reliant on the characteristics of larger individuals.

Overall, differences between the transient dynamics of the tropical and subtropical *Acropora* spp. populations were underpinned by variation in the vital rate patterns of existing colonies. However, recruitment is a fundamental component in the dynamics and resilience of coral communities (Adjeroud *et al*. 2017). Accordingly, we observed that recruitment patterns actually exerted the largest absolute influence on the transient dynamics of the two populations, although, this influence was masked by contrasting patterns in larval settlement and recruit survival (Fig. 3A). We observed that the settlement of *Acropora* larvae was lower in the subtropics compared with the tropics. With abiotic barriers limiting the dispersal and survival of coral larvae at higher latitudes this pattern is to be expected (Nakabayashi *et al*. 2019), despite conflicting recent evidence of warming induced increases in the densities of settling subtropical larvae (Price *et al*. 2019). Intriguingly though, we report that the survival of coral larvae following successful settlement appeared highest in the subtropics. Whilst consistent with expected density dependant patterns in the survival of newly settled larvae (Cameron & Harrison 2020), our finding disagrees with previous reports of extremely high annual post-settlement larval mortality within a subtropical environment (Wilson & Harrison 2005). Seawater temperatures at the time of settlement influence the survival of coral larvae (Randall & Szmant 2009). Equally, acroporid corals are known to be highly sensitive to cold shock (short-term exposure to cold temperatures; Roth *et al*. 2012). Therefore, with our assessment of recruitment patterns reliant on settlement plates and plot surveys occurring during boreal summer months we acknowledge that our estimates of subtropical recruit survival may represent overestimates arising from the inclusion of individuals yet to experience the selective pressures of cooler subtropical seasons.

## Conclusions

Understanding the extent to which marginal populations can embrace environmental variation, and the mechanisms that underpin the success, or failure, of populations along range boundaries, is necessary if we are to anticipate the continued viability of crucial species, communities, and ecosystems (Valladares *et al*. 2014; Merow *et al*. 2017). Equally, distinguishing how vital rate characteristics manifest under differing environmental regimes will help resolve the climate envelopes of different species and ecosystems, allowing for more accurate predictions of population persistence or collapse (Trisos *et al*. 2020). Climatic warming is facilitating the poleward expansion of coral populations into subtropical coastal ecosystems (Beger *et al*. 2014; Vergés *et al*. 2019). The dynamics of coral populations establishing within tropicalising environments offer valuable insights into the ability of coral communities for persisting within suboptimal habitats and adapting to future, more variable, climates (Camp *et al*. 2018). However, our lack of an appreciation for the demographic characteristics of coral populations and their abiotic drivers (Edmunds *et al*. 2014; Edmunds & Riegl 2020), inhibits our capacity for exploring these insights.

The transient dynamics of populations define their responses to disturbance, and ultimately their dynamics within variable environments (Hastings 2004; Stott *et al*. 2011; McDonald *et al*. 2016; Hastings *et al*. 2018). Transient demographic theory has advanced our understanding of invasive potential, allowing us to forecast the ability of species to establish populations outside their core range (Iles *et al*. 2016; Merow *et al*. 2017; Jelbert *et al*. 2019). We have illustrated here how the transient dynamics of coral populations coordinate their establishment at higher latitudes, mediating their response to enhanced seasonal variation. Equally, *Acropora* spp. populations in southern Japan, display the demographic plasticity necessary for the continued exploitation of higher latitude environments. However, with this work we have only begun to gather evidence of the mechanisms supporting the redistribution of coral populations. It is crucial we continue evaluating how patterns in the transient dynamics of coral populations translate across various species, and over broader spatial scales. Without improving current knowledge regarding the dynamics of coral populations we will be unable to predict the persistence and future reassembly of coral communities and their associated reef taxa (Edmunds & Riegl 2020; Pisapia *et al*. 2020; Cant *et al*. 2021a).

## Acknowledgements

The authors would like to thank I. Mizukami, H. Kise, C. Fourreau, G. Masucci, P. Biondi (all University of the Ryukyus), S. Nishihira (Dive Team Snack Snufkin), M. Tamae, H. Nakakoji (both Marine Space), and all staff at both Pacific Marine and SeaAir for help in field data collection. Funding for this research was provided by a Natural Environment Research Council (NERC) Doctoral Training Programme Scholarship to JC, grants by the British Ecological Society, UK, the Winifred Violet Scott Estate, Australia, and the European Union’s Horizon 2020 research and innovation programme under the Marie Skłodowska-Curie grant agreement TRIM-DLV-747102 to MB, and ORCHIDS project funding from the University of Ryukyus to JDR. Recruitment surveys were supported by JSPS KAKENHI Grant Number 16K07527 to MN. RS-G was supported by a NERC Independent Research Grant (NE/M018458/1).

