## Supplementary material for "Transient amplification enhances the persistence of tropicalising coral populations in marginal high latitude environments"

**S1. Quantifying the variation between the patterns of colony survival, growth, and fragmentation within tropical and subtropical environments.**

To explore the influence of environmental variability on the transient dynamics of coral populations, we constructed Integral Projection Models (IPMs; Easterling *et al.* 2000) describing the respective dynamics of *Acropora* spp. populations from a tropical and subtropical environment. We parameterised our IPMs through an annual census of *Acropora* spp. colonies conducted in southern Japan between 2017 – 2019 (Fig. 1). In September 2017, we tagged *Acropora* individuals within the tropical reef communities of Okinawa at Hentona (26.75˚, 128.18˚), Oura Bay (26.54˚, 128.08˚), and Miyagi Channel (26.35˚, 127.99˚), using permanently marked plots (n = 32 plots, 2.4 ± 0.19 [SD] colonies/plot). Permanent plots were also set up within the subtropical communities of Kochi, Shikoku, to tag *Acropora* colonies at Okinoshima (32.75˚, 132.55˚), Kashiwajima (32.77˚, 132.62˚), and Nishidomari (32.78˚, E 132.73˚; n = 35 plots, 6.2 ± 0.25 colonies/plot). We assembled these permanent plots by fixing numbered tags into bare reef substrate, with each plot consisting of a tag and the surrounding coral colonies within a 2m^2^ area. Photographs, with scale bars included for reference, were used to capture the visible horizontal extent of all *Acropora* colonies within each plot. We estimated the horizontal surface area (cm^2^) of tagged colonies using ImageJ (Schneider *et al.* 2012), before log-transforming colony sizes prior to further analyses to normalise the size distribution and improve the resolution of smaller colonies.

*Colony survival*

Repeated surveys of tagged colonies in August 2018, and August 2019, allowed us to quantify size-specific patterns in colony survival. Colony survival was recorded in the field as the presence or absence of tagged colonies during successive surveys, and modelled as a function of colony size at time *t* using a logistic regression (Fig. S1). When modelling this relationship we included the fixed effect of region (tropical or subtropical) allowing us to compare survival patterns across our tropical and subtropical populations. To parameterise patterns in colony survival, we used colony data pooled across years to ensure greater statistical power within our analyses. Our regression models for this vital rate therefore included colony identity as a random variable to account for repeated measures of individual colonies. We also initially included survey site location as a random effect to account for any nesting within our data, although this resulted in a singular fit, due to insufficient data to support the subsequent complexity of the model. Thus, we excluded the random effect of site to prevent overfitting within our model allowing us to best explore trends within the data. Overall, the probability of survival increased with colony size in both populations, although this trend was more pronounced within the subtropical population (Fig. S1).

**Figure S1.** Size specific patterns in the survival probabilities of *Acropora* spp. colonies within a tropical (Red) and subtropical (Blue) setting. Shaded regions represent 95% Confidence Intervals.

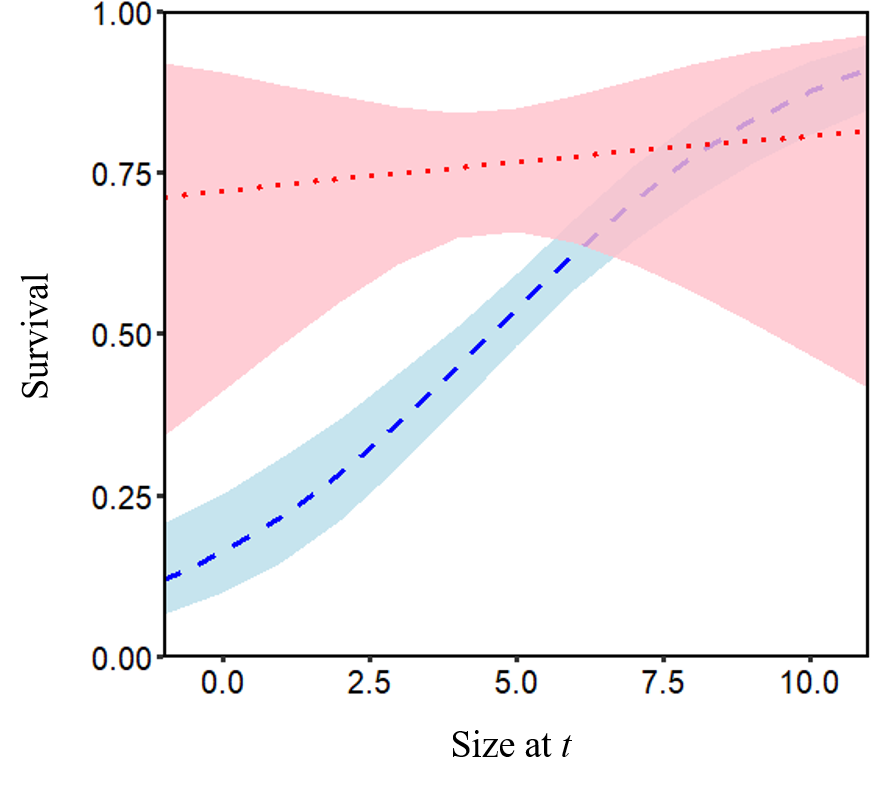

*Colony size distributions & changes in colony size*

Revisiting tagged *Acropora* colonies in both Okinawa and Kochi allowed us to explore temporal trends in the size distributions of the tropical and subtropical populations (Fig. S2). We observed that between 2017 and 2019, both populations displayed declines in the dominance of larger individuals, and that in any given year, mean colony size was largest in the subtropics (Table S1). However, trends in the skewness of each population’s annual size distribution indicated transitions toward greater righthand skew, reflecting an increase in the density of smaller colonies; a trend that was particularly true for the subtropical population during 2019 (Table S1).

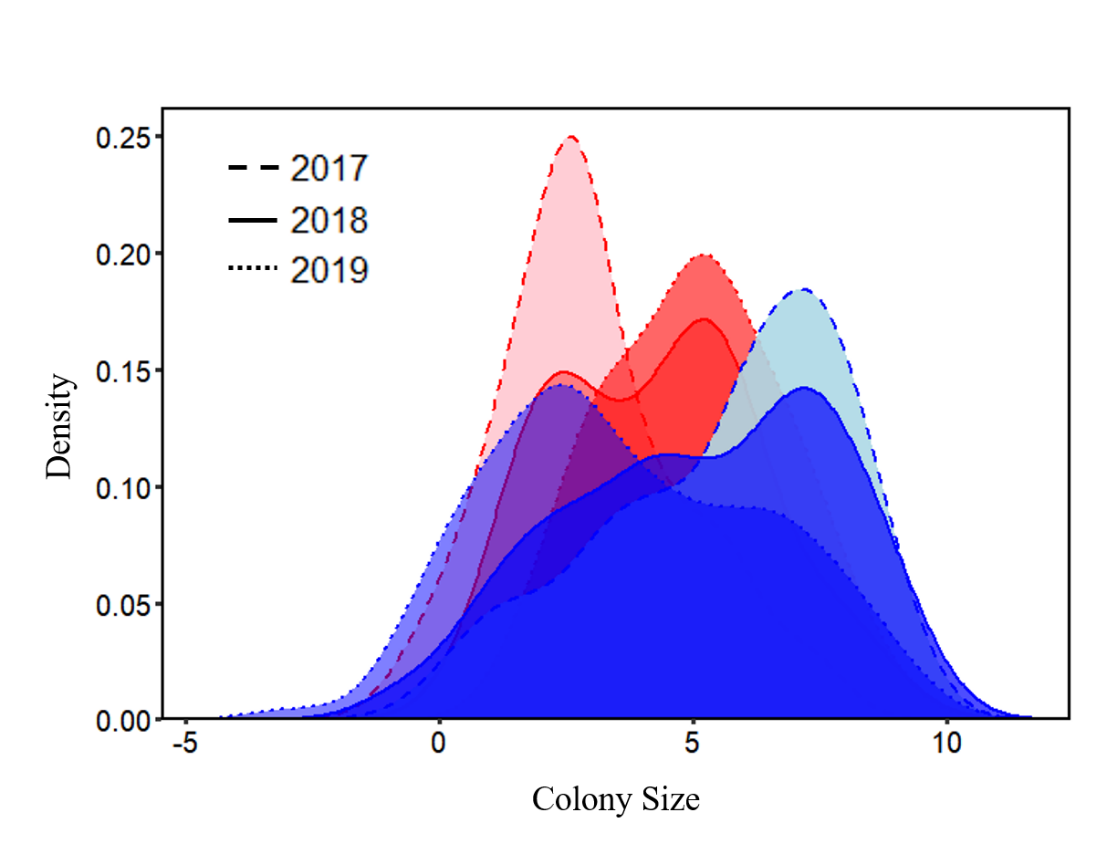

**Figure S2.** Temporal trends in the size distributions of tagged tropical (Red) and subtropical (Blue) *Acropora* spp. populations recorded during annual surveys between 2017 and 2019.

**Table S1.** Temporal trends in mean colony size and skew in the size distributions of tagged tropical and subtropical *Acropora* spp. populations recorded during annual surveys between 2017 and 2019. Error reported as 95% CI.

|  | **Subtropical** | | | **Tropical** | |
| --- | --- | --- | --- | --- | --- |
| Census | MEAN | SKEW | MEAN | | SKEW |
| 2017 | 5.62 [5.27, 5.96] | -0.63 | 5.00 [4.39, 5.62] | | 0.05 |
| 2018 | 5.14 [4.77, 5.52] | -0.30 | 4.27 [3.67, 4.88] | | 0.19 |
| 2019 | 3.69 [3.35, 4.04] | 0.18 | 2.99 [2.56, 3.42] | | 0.48 |

We also quantified size-specific transitions in colony size using the difference between colony surface areas recorded during successive annual surveys. Patterns in colony size transitions were calculated using linear regression and reflected the relationship between colony size at time *t* and size at time *t+1.* As with our models of colony survival, we modelled colony growth with region included as a fixed effect, and colony identity included as a random variable, but excluded site as a random effect to avoid overfitting our models. Again, data regarding the annual size transitions of tagged colonies was pooled across years to enhance the resolution of our analyses. We found that patterns in the size transitions of colonies were largely consistent across both the tropical and subtropical populations, with enhanced positive growth reported in smaller colonies and stasis in larger colonies (Fig. S3). We also separately modelled the relationship between the variance in colony size at time *t+1* and colony size at time *t*. This relationship was determined by modelling the residuals from our colony growth model above, as a function of colony size at time *t.* For this relationship we assumed a gamma distribution, allowing for a non-linear pattern whilst preventing negative
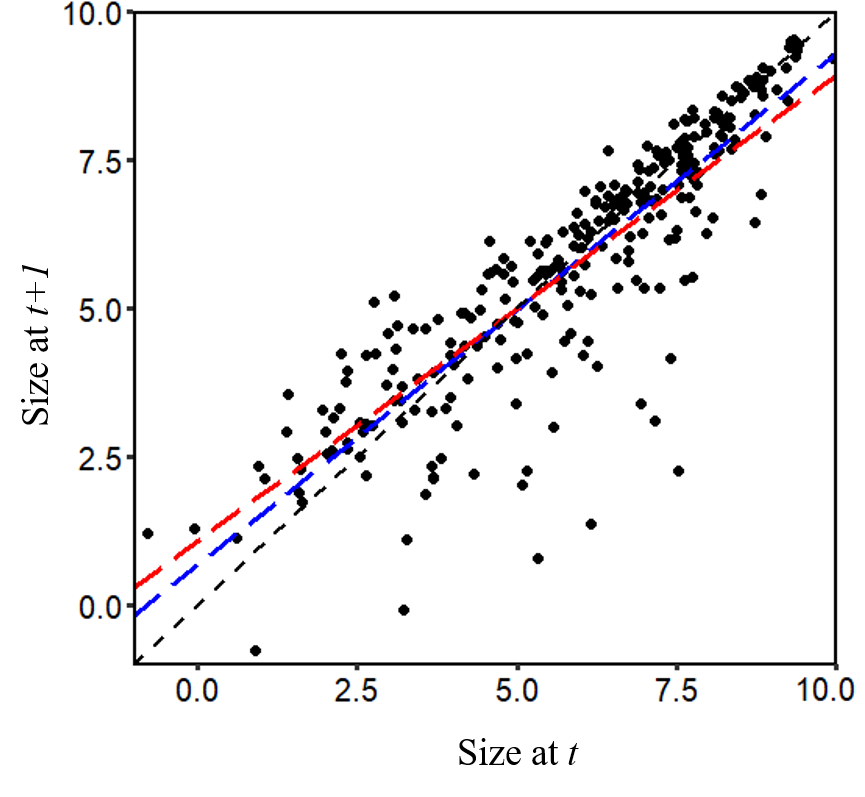
variance. For clarity, we did compare this gamma model with a linear regression format, however, model AIC scores confirmed our assumption of a gamma distribution to be appropriate (AIC: linear = 572.6; gamma = 326.5).

**Figure S3.** Size specific patterns in the size transitions of *Acropora* spp. colonies within a tropical (Red) and subtropical (Blue) setting. 1:1 diagonal line represents no change in size between times *t* and *t+1*.

*Colony fragmentation*

Through the demographic census of tagged colonies we also estimated size-specific patterns in the probability of colony fragmentation. During each survey, we reported if tagged colonies had fragmented following observed evidence of colony breakage, but only in the event that colony remnants remained visible in order to distinguish between fragmentation and partial mortality. We modelled the probability of colony fragmentation during the interval *t* to *t+1* as a function of colony size at *t* using a logistic regression. However, due to the low frequency of annual fragmentation events (number of events reported, $n_{i}^{\kappa}$) within our tropical population during both 2018 ($n_{tropical}^{\kappa}$ = 1, $n_{subtropical}^{\kappa}$ = 15, $n_{total}^{\kappa}$= 16), and 2019 ($n_{tropical}^{\kappa}$ = 5, $n_{subtropical}^{\kappa}$ = 14, $n_{total}^{\kappa}$= 19), we explored patterns in the probability of fragmentation using data pooled from both the tropical and subtropical populations (Fig. S4A). We modelled the vital rate of fragmentation without the random effect of colony identity since fragmentation did not occur repeatedly enough across individuals to support the inclusion of this random variable. To ensure our approach of pooling data across populations did not prevent us from suitably capturing the differing prevalence of fragmentation within the dynamics of our tropical and subtropical populations, fragmentation patterns across our models were subsequently weighted according to the relative proportion of annual events recorded in our tropical and subtropical plots (${n_{i}^{\kappa}}/{n_{total}^{\kappa}}$). Finally, alongside each reported fragmentation event, the quantity and size (surface area, cm^2^) of all colony fragments produced were also recorded with, in each case, the largest fragment retaining the original colony’s identity. We subsequently used these data to estimate patterns in both fragment production and fragment size at time *t+1*, as a function of initial colony size at time *t*, using linear regression (Fig. S4B & C).

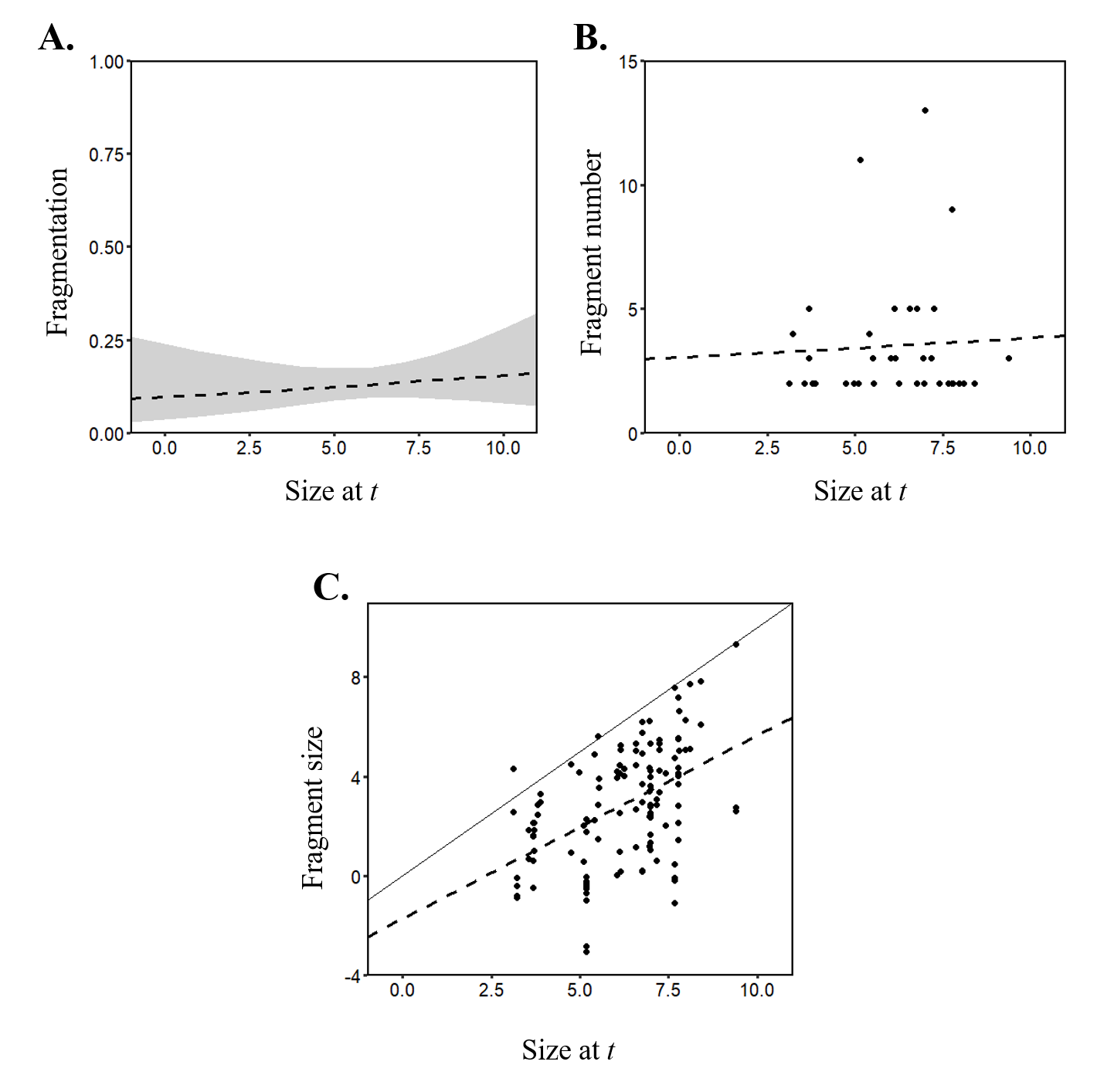

**Figure S4.** Size specific patterns in (**A**) the probability of fragmentation, (**B**) the number of fragments produced, and (**C**) the size of subsequent fragments, observed in tagged *Acropora* spp. colonies within southern Japan. Solid line in panel C represents the initial size of colonies prior to fragmentation.

**S2. Evaluating patterns of recruitment within tropical and subtropical coral populations.**

Across our IPMs recruitment was represented using the three vital rates of colony fecundity (*f_n_*), the probability of larval settlement (*ψ*), and the subsequent probability of recruit survival (*Ϯ*). We did not directly measure colony fecundity using the tagged colonies. Instead we sourced data on the size-specific fecundity (larval volume, cm^3^) of *Acropora* colonies on the Great Barrier Reef (Hall & Hughes 1996) from the Coral Trait Database (Madin *et al.* 2016). These data describe an exponential increase in the larval output of *Acropora* colonies with increasing colony size, and subsequently, using non-linear least squares regression, we applied this relationship to estimate the expected larval output of each tagged colony, given their size in each year. Crucially, quantifying fecundity in this way assumes that the reproductive output of corals remains fixed across varying environments. However, this is a reasonable assumption given that the reproductive biology of corals is consistent across monophyletic clades, families, genera, and species (Baird *et al.* 2009; Harrison 2011). We have also addressed this assumption within our demographic models through the inclusion of the larval settlement and recruit survival parameters (see below) which constrained recruitment patterns within our models according to empirical observations made within the tagged populations. Equally, our use of the larval settlement parameter also served to translate colony fecundity estimates from measures of larval volume into expected counts of settling larvae within tropical and subtropical environments.

Alternatively, we estimated the parameters of larval settlement probability (*ψ*) and recruit survival probability (*Ϯ*), for the tropical and subtropical populations, using larval counts made during prior settlement tile surveys conducted in both Okinawa (Nakamura *et al.* 2015) and Kochi (Nakamura, *unpublished data*). Between 2011 and 2013, Nakamura *et al.* (2015) annually deployed sets of settlement plates at seven sites along the Onna coastline of Okinawa (Fig. 1A), to quantify spatial and temporal variation in the number of settling *Acropora* larvae. Each plate set consisted of two 10×10cm cement tiles fastened one above the other separated by a 2cm gap. Following pre-conditioning each plate set was deployed on the reef over a two month period coinciding with local *Acropora* spawning events. Upon retrieval, across each tile set, only the upper and lower surfaces of the top tile and the upper surface of the lower tile were observed for coral spat, with the lower surface of the lowest tile having been used to secure both tiles to the reef. Subsequently, each plate set reflected a settlement area of 300cm^2^. A similar method was applied in Kochi, at Kashiwajima and Nishidomari during 2016–18 (Fig. 1A), providing a comparison between the annual rates of larvae settlement for *Acropora* spp. in both a tropical and subtropical setting. Again plate sets consisting of two 10x10cm cement tiles were deployed at Kashiwajima and Nishidomari. However, a key difference on this occasion was that upon retrieval of the plate sets both surfaces of both tiles were observed for coral spat. Thus, for these subtropical counts, each plate set reflected a settlement area of 400cm^2^.

We used these settlement counts to estimate the mean number of settling larvae per plate set which we subsequently scaled up to reflect larval settlement per unit area at the spatial scale of our plots (~2m^2^; Table S2). Dividing the total estimated larval outputs for our tagged tropical and subtropical *Acropora* colonies during each annual interval (2017-18 & 2018-19), by the corresponding regional average count of larval settlement per unit area, we were then able to determine ratios translating colony fecundity into the expected number of settling larvae (*ψ*; *sensu* Bramanti *et al.* 2015) within tropical and subtropical environments (Table S2). Next we used our scaled estimates of larval density, and empirical counts of new *Acropora* colonies appearing within our tagged plots each year, to quantify ratios describing the annual post-settlement survival probability within both tropical and subtropical settings (*Ϯ*; Table S2).

| **Table S2.** Temporal trends in larval settlement, and recruit survival within *Acropora* spp. populations in Okinawa and Kochi. Scaled larval densities were estimated by extrapolating mean settlement tile counts to reflect the spatial coverage of our tagged plots (~2m^2^). The scaled larval densities were then combined with estimates of total colony fecundity and empirical recruit counts from 2017-18 and 2018-19, to determine annual estimates of larval settlement probability (*ψ*) and recruit survival probability (*Ϯ*). Error displayed as 95% CI. | | | | | | |
| --- | --- | --- | --- | --- | --- | --- |
|  | Scaled larval density  (larvae plot^-1^) | Larval settlement probability (*ψ*) | | Recruit density  (Recruit plot^-1^) | Recruit survival probability (*Ϯ*) | |
| OKINAWA | 1175.26  [1135.87, 1214.65] | 2017-18 | 0.2143  [0.2071, 0.2215] | 10 | | 0.0085  [0.0082, 0.0088] |
|  |  | 2018-19 | 0.2997  [0.2896, 0.3097] | 28 | | 0.0238  [0.0231, 0.0247] |
| KOCHI | 31.11  [22.96, 39.26] | 2017-18 | 0.0004  [0.0003, 0.0005] | 20 | | 0.6429  [0.5094, 0.8710] |
|  |  | 2018-19 | 0.0005  [0.0004, 0.0006] | 63 | | 2.0250  [1.6047, 2.7436] |

We acknowledge, here, that our approach to implement scaled settlement tile counts in estimating recruitment parameters entails two important considerations. Firstly, coral larvae predominantly settle close to the edge of settlement tiles resulting in a potential underestimation of larval settlement per unit area when scaling any counts (Price *et al.* 2019). Secondly, coral larvae are selective with settlement surfaces (Norström *et al.* 2007; Arnold *et al.* 2010) and identify suitable locations through a complex series of biotic cues (Price 2010). Accordingly, the area of substrate represented by our tagged plots may not be equivalent to the area of available effective substrate for larval settlement, leading to overestimates when scaling tile counts. Despite these pitfalls, scaling tile counts to more representative dimensions remains a common technique within recruitment assessments (Price *et al.* 2019), and with both these features opposing each other with regards to their impact on our capacity to accurately model regional recruitment patterns, we did not explicitly account for them within our parameter estimation.

Finally, during plot surveys in 2018 and 2019 we recorded the size (cm^2^) of new colonies appearing within the plots to provide a measure of recruit size (*C_0_*) and how this varies between tropical and subtropical populations (Fig. S5). Due to the difficulties associated with identifying the parental lineage of new recruits, recruit size was modelled independently of parent colony size using linear regression. This approach is consistent with evidence that the complex dynamics of larval settlement and survival are strictly coordinated by synergistic abiotic constraints (Vermeij *et al.* 2009; Doropoulos *et al.* 2016), and therefore independent from parental characteristics. Initially, we modelled recruit size with ecoregion (subtropical *vs.* tropical) and survey year, included as fixed effects. However this approach demonstrated no significant difference between the recruit size distributions of the tropical and subtropical populations, and little within-population variation between years (Fig. S5; GLM: F_3,117_ = 1.09, *p = 0.36*). We subsequently, dropped the term of survey year from our model of recruit size, but retained ecoregion as a fixed effect as AIC scores confirmed this to be the most appropriate model fit (AIC: no terms = 383.8; ecoregion only = 383.2, ecoregion * year = 386.5).

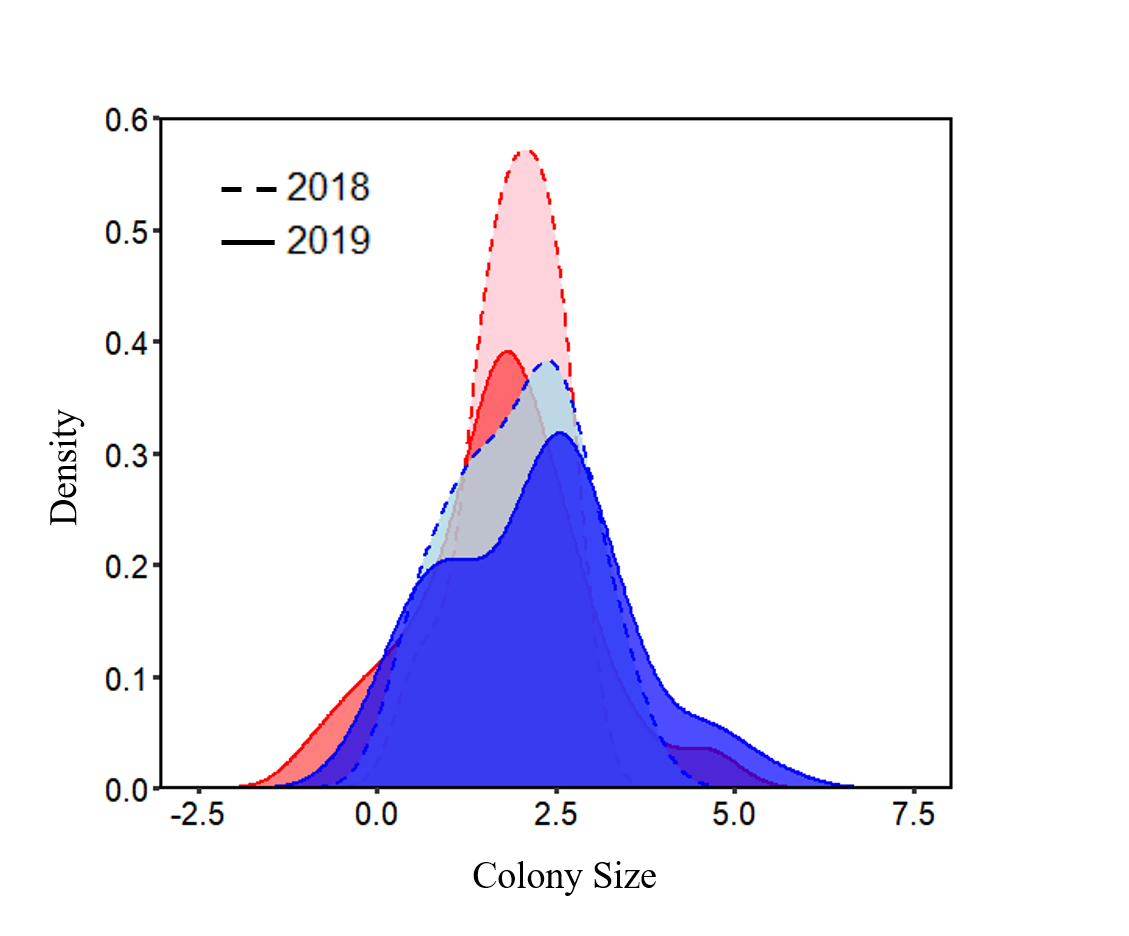

**Figure S5.** Temporal trends in the size distributions of surviving recruit cohorts observed within tropical (Red) and subtropical (Blue) *Acropora* populations during 2018 and 2019.

**S3. Exploring latitudinal trends in the traits of *Acropora* spp. populations.**

We have demonstrated that there exists a latitudinal trade-off between the long-term persistence and short-term exploitation potential in *Acropora* spp. populations. In the subtropics populations of *Acropora* spp. display a greater capacity for demographic compensation following disturbances, whereas in the tropics populations instead exhibit more stable asymptotic growth rates (*λ*). We admit, however, that any interpretations of our results need to be approached with a degree of caution. Due to unresolved coral taxonomies and the prevalence of numerous cryptic species, it is difficult to identify *Acropora* spp. individuals to the species level in the field (Fukami *et al.* 2004; Richards & Hobbs 2015; Richards *et al.* 2016). Thus we conducted all our analyses at the genus level. Yet without explicit data regarding the species compositions of our tagged tropical and subtropical *Acropora* populations it would be inappropriate for us to ignore the fact that our reported demographic trends may simply coincide with latitudinal shifts in species configurations, rather than any demographic plasticity. Subsequently, it was necessary to disentangle the potential role shifts in species compositions, and therefore shifts in species traits between tropical and subtropical environments, play in mediating the demographic variation we observed.

Using the Coral Trait database (Madin *et al.* 2016) we extracted trait values for a series of numerical and categorical traits for *Acropora* species known to occur within the coral communities of Okinawa and/or Kochi (Table S3). As expected, species richness was highest in the tropics with 73 different species of *Acropora* reported in Okinawa. Alternatively, just 26 species have been recorded within the *Acropora* spp. communities of Kochi, of which only four are not found in Okinawa (Table S3). To explore variation in the characteristics between these *Acropora* spp. communities in southern Japan, we focused on seven numerical traits relating to calcification rate (µm cm^-2^ h^-1^), corallite width (maximum and minimum; mm), depth (maximum, minimum, and mean; m), and colony growth rate (mm year^-1^), alongside the three categorical traits of growth form, water clarity preference, and wave exposure preference. A single value for each trait was extracted to describe the characteristics of each *Acropora* species. For any species for which multiple estimates had been reported for any given trait we retained the mean (numerical traits), or modal trait value (categorical traits).

We used the extracted trait values to quantify how the characteristics of *Acropora* spp. communities vary between tropical and subtropical environments (Table S4; Fig. S6). T-tests confirmed that there exists no significant variation between the distributions of any of the selected numerical traits, between the tropical and subtropical communities (Table S4). Equally, the proportional arrangements of the selected categorical traits remain similar across the communities from the two regions (Fig. S6). An exception to this trend is that there is a change in the composition of colony growth forms between the two regions. In the tropics there is a greater prevalence of arborescent morphologies (Fig. S6C). However, in the subtropics there is a shift towards increased exploitation of tabular growth forms (Fig. S6C), consistent with the need for *Acropora* spp. individuals to maximise their ability to compete for access to photosynthetic radiation, which is more limited at higher latitudes (Muir *et al.* 2015; Zawada *et al.* 2019).

Overall the trait characteristics of *Acropora* species associated with tropical and subtropical environments appears fixed across both regions; providing evidence that our reported trade-off between demographic compensation and demographic stability is not a consequence of a shift in the species composition of tropical and subtropical *Acropora* spp. communities in southern Japan. Moreover, given that there is considerable overlap between the species configurations of the *Acropora* spp. communities of Okinawa and Kochi we can be more confident that our findings do indeed present evidence of demographic plasticity associated with the need for coral populations in subtropical environments to enhance their viability despite increased environmental variability.

**Table S3.** List of *Acropora* species reported by Nishihira & Veron (1995) and Veron *et al.* (2016) to occur within the coral communities of Okinawa and/or Kochi. Colour to the left of each species reflects its recorded distribution: Okinawa only (Red), both Okinawa & Kochi (Orange), and Kochi only (Blue).

|  | *A. abrolhosensis* |  | *A. longicyathus* |  | *A. anthocercis* |
| --- | --- | --- | --- | --- | --- |
|  | *A. abrotanoides* |  | *A. microclados* |  | *A. aspera* |
|  | *A. aculeus* |  | *A. microphthalma* |  | *A. copiosa* |
|  | *A. acuminata* |  | *A. millepora* |  | *A. cuneata* |
|  | *A. akajimensis* |  | *A. monticulosa* |  | *A. dendrum* |
|  | *A. austera* |  | *A. nana* |  | *A. divaricata* |
|  | *A. awi* |  | *A. nobilis* |  | *A. florida* |
|  | *A. brueggemanni* |  | *A. palifera* |  | *A. hyacinthus* |
|  | *A. carduus* |  | *A. paniculata* |  | *A. insignis* |
|  | *A. cerealis* |  | *A. parilis* |  | *A. latistella* |
|  | *A. clathrata* |  | *A. pichoni* |  | *A. listeri* |
|  | *A. cytherea* |  | *A. pulchra* |  | *A. loripes* |
|  | *A. danai* |  | *A. robusta* |  | *A. nasuta* |
|  | *A. digitifera* |  | *A. rosaria* |  | *A. samoensis* |
|  | *A. echinata* |  | *A. sarmentosa* |  | *A. solitaryensis* |
|  | *A. efflorescens* |  | *A. secale* |  | *A. striata* |
|  | *A. exquisita* |  | *A. sekiseiensis* |  | *A. subulata* |
|  | *A. formosa* |  | *A. selago* |  | *A. teres* |
|  | *A. gemmifera* |  | *A. subglabra* |  | *A. tumida* |
|  | *A. grandis* |  | *A. tenella* |  | *A. valida* |
|  | *A. granulosa* |  | *A. tenuis* |  | *A. verweyi* |
|  | *A. horrida* |  | *A. valenciennesi* |  | *A. willisae* |
|  | *A. humilis* |  | *A. vaughani* |  | *A. glauca* |
|  | *A. inermis* |  | *A. wallaceae* |  | *A. japonica* |
|  | *A. irregularis* |  | *A. yongei* |  | *A. pruinosa* |
|  | *A. kirstyae* |  |  |  | *A. stoddarti* |

| **Table S4.** Comparison of the numerical trait characteristics of tropical and subtropical *Acropora* spp. communities from Okinawa and Kochi, respectively. T-tests were used to evaluate for any statistical significance in the trait distributions from the two regions. Error reported as ±SD. | | | | |
| --- | --- | --- | --- | --- |
| Trait | | Okinawa | Kochi | Test statistic |
| CALCIFICATION RATE  (µm cm^-2^ h^-1^) | | 1.26 ± 1.18 | 1.35 ± 1.31 | *p* = 0.90 |
| CORALLITTE WIDTH (mm) | Max | 1.20 ± 0.29 | 1.24 ± 0.26 | *p =* 0.64 |
|  | Min | 0.62 ± 0.20 | 0.69 ± 0.25 | *p =* 0.30 |
| DEPTH  (m) | Max | 25.08 ± 9.57 | 22.59 ± 8.32 | *p* = 0.26 |
|  | Mean | 14.82 ± 6.53 | 12.82 ± 4.75 | *p* = 0.14 |
|  | Min | 4.56 ± 4.75 | 3.04 ± 2.38 | *p* = 0.06 |
| GROWTH RATE  (mm year^-1^) | | 50.50 ± 44.96 | 35.62 ± 21.86 | *p* = 0.28 |

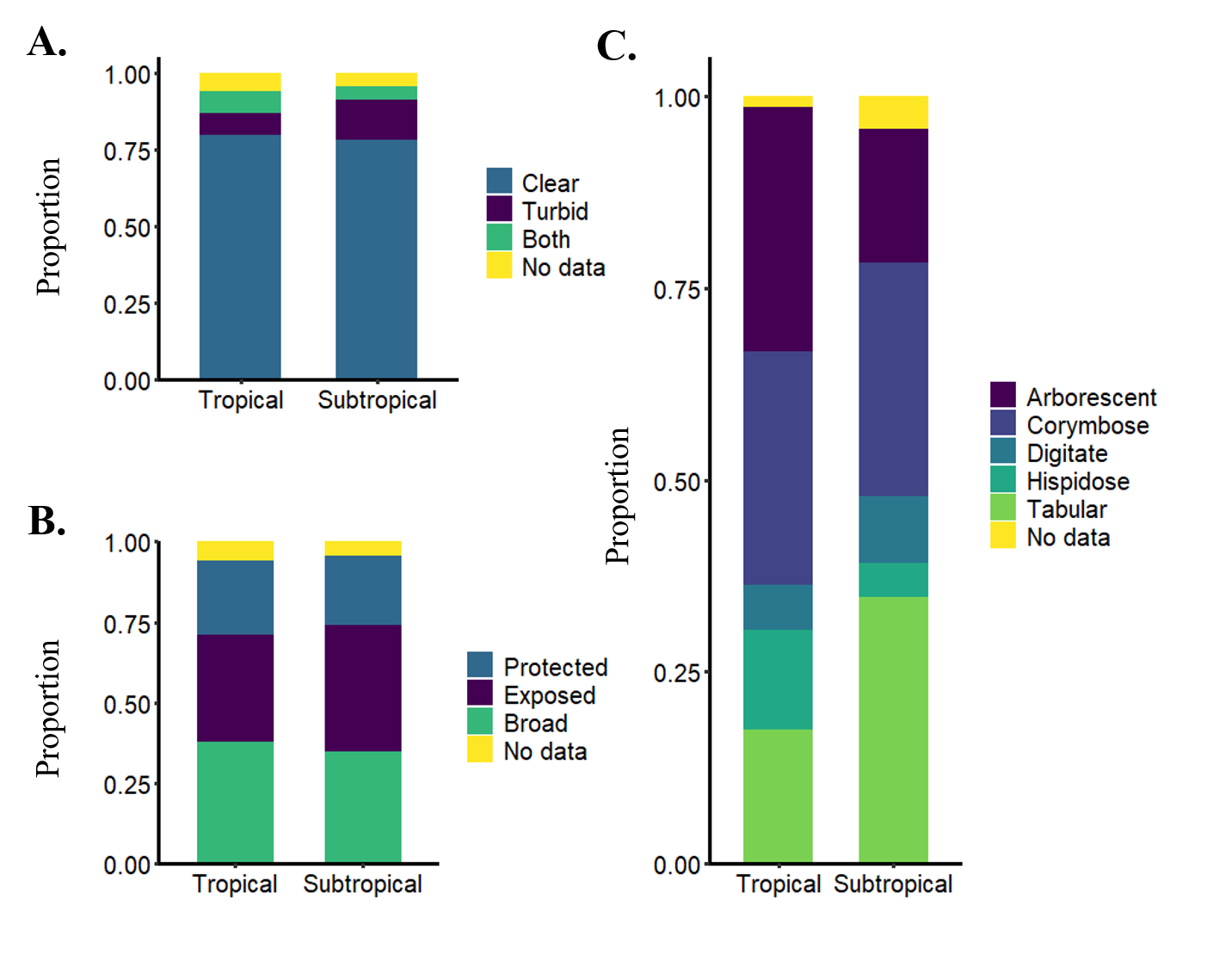

**Figure S6.** Patterns in the **categorical traits of (A)** water clarity preference, **(B)** wave exposure preference and **(C)** growth form within the respective tropical and subtropical *Acropora* spp. communities of Okinawa and Kochi. Proportions show the number of species exhibiting each trait relative to the number of species present in each region (Tropics, n = 73; Subtropics, n = 26).
